# The reproductive status of the host determines the tolerance and resistance to *Mycobacterium marinum* infection in *Drosophila melanogaster*

**DOI:** 10.1101/2022.08.30.505810

**Authors:** Marta Arch, Maria Vidal, Esther Fuentes, Anmaw Shite Abat, Pere Joan Cardona

## Abstract

Both the sex and the reproductive status of the host have a major impact on the regulation of the immune response against infection. Due that *Drosophila melanogaster* has become a powerful model for study such interactions, we wanted to understand whether the sex or the reproductive status has an impact on the tolerance or resistance of the host to the in the model of systemic *Mycobacterium marinum* infection. We measured gene expression by RT-qPCR of immune genes, *diptericin* and *drosomycin*, as well as host survival and the bacillary load at time of death. We also assessed the impact of metabolic (by expression of *upd3* and *impl2*) and hormonal (by *ecR* expression) regulation in the protection against infection. Data showed that resistance increased in actively mating males and females, and also in mated females while reducing the tolerance to infection. The data also suggest the Toll and IMD pathways determine tolerance and resistance, respectively, while basal levels of *ecR* favors the stimulation of the IMD pathway. A time related role has been found for *upd3* expression, linked to the increase or decrease in the mycobacterial load at the beginning and at the end of the infection, respectively. Finally, *impl2* expression has been related to the increase of resistance only in non-actively mating males. The results obtained allows further interpretation of the results when assessing the differences between sexes and highlights the role of the reproductive status in *D. melanogaster* to face infections, since our data demonstrated their importance to determine resistance and tolerance against *M. marinum* infection.

## 1. Introduction

Reproduction and immunity are resource-intensive traits in their deployment and maintenance, thus, a trade-off exists between immunity and reproduction when the resources are limited [1]. This trade-off has been documented in both mammals and non-mammals [2–5]. Its study is important to shed light on the impact of directional selection and evolution in the maintenance or the variation of these traits, thus helping to better characterize the relationship between these two physiological mechanisms [6].

In this regard, *Drosophila melanogaster* is a powerful model organism to study the interaction between reproduction and immunity due to its already well-studied courtship ritual and its well-characterized immune system [7]. Immunity in *D. melanogaster* relies solely on innate immune responses, nevertheless, there are a lot of aspects of this innate immunity that are conserved between flies and mammals, such as the organization of the immune response into humoral and cellular arms, and the presence of NF-κB family transcription factors and signal transduction pathways [8]. The humoral response is mainly based on the synthesis of antimicrobial peptides (AMPs) in the fat body of the fly. This process is regulated by two signalling pathways: the Toll and the immune deficiency (Imd) pathways, which activate different NF-κB-like factors and induce the production of Drosomycin and Diptericin (as the main AMP of each pathway, respectively) [9,10]. Flies also have a simpler but complete JAK/STAT pathway, which regulates a wide range of biological processes in the fruit fly, including immunity [11,12]. Regarding cellular immunity in *Drosophila*, it is mediated by hemocytes. When a septic injury, tissue damage, or exposure to lipopolysaccharides (LPS) occurs, hemocytes trigger the expression of Unpaired (Upd) 3, which activates the JAK/STAT pathway in the fat body, which in turn contributes to the humoral response development and leads to upregulation of immune genes [11,13].

It is well known that hormones and metabolism, as well as circadian cycles, have central roles in the regulation of systemic physiology, including immunity, in both insects and mammals [14–16]. The metabolic homeostasis in insects is mainly regulated by the insulin/insulin-like signalling (IIS) pathway. In *D. melanogaster* the activation of this pathway leads to the storage of energy in form of triglycerides in the fat body. Interestingly, after an immune stimulus, the immune pathways interact with the IIS pathway in the fat body resulting in loss of energy storage and suppression of host growth [17–19]. This fact has been recently described as the “selfish immune system theory”, which is based on the fact that activated phagocytes release signalling molecules (selfish immune factors, SIFs) that regulate host energy to steal resources from other non-immune tissues to induce an efficient acute immune response, thus inhibiting insulin signalling during infection [20–23]. Authors propose the insulin/IGF antagonist Imaginal morphogenesis protein late 2 (Impl2) – which has been identified as a cancer-derived cachectic factor in flies –, and Upd3 as potential SIFs in *Drosophila* [24–26].

Hormones also play an important role in the regulation of immunity [14]. Both in mammals and insects exists a sexual dimorphism affecting immunity (survival, pathology, bacterial load and activity in response to infections) at basal conditions, after a pathogenic challenge, and even upon ageing [27,28]. Few studies about the topic revealed that these differences might be mediated by different immune players depending on the pathogen and the route of infection. For example, males appeared to be more resistant to systemic infections by *Providentia* species, and *Enterococcus faecalis*, while females seemed to be more resistant to *Pseudomonas aeruginosa, Staphylococcus aureus* and *Serratia* infections [29–31]. Recently, a role for the Toll pathway in mediating sex dimorphisms has arisen with the finding that the loss of Toll-7 reduces resistance to *P. aeruginosa* and *E. faecalis* in males but not in females [32]. In mammals, steroid hormones have been linked with these sex differences, and interactions between these type of hormones and the innate immune system of *Drosophila* have also been described [33,34]. The steroid hormone Ecdysone is the main regulator of the insect life cycle [35]. In adult flies, Ecdysone is produced in the ovaries after mating, thus showing higher levels in females than in males [36–38]. Its signalling through the receptor complex formed by the Ecdysone receptor (EcR) and Ultrapiracle (Usp) is required for the proper expression of the pathogen-sensing receptor PGRP-LC and, thus, the production of Imd-dependent AMPs [39] and the cellular immunity [34,40]. At the same time, Ecdysone levels are regulated by stress signals and it has been related to age-related immune changes, as its depletion increases *Drosophila’s* lifespan [41,42].

Nevertheless, it is not only sex that affects immunity, but also reproductive status. In *Drosophila*, mated females showed decreased survival, higher pathogen loads and reduced AMPs production after pathogenic infections [43–45], although mating does not affect the clearance of non-pathogenic *E.coli* [46,47], thus making the trade-off between mating and immunity evident only when the infection is pathogenic [4]. On the contrary, increased resistance to infections correlates with a reduction in egg viability [48], but female flies unable to generate eggs do not have reduced immunity after mating [45] and females fed with yeast *ad libitum* improved both fecundity and resistance to infections proving that reproduction and immunity also compete for energy resources [49]. Along the same lines, males exposed to a higher number of females also showed increased susceptibility to bacterial infections [50]. Conversely, it has been also demonstrated that, despite the significant metabolic cost of infection, flies don’t change and even increase the ability to survive infection and clear it [7,51]. Likewise, mated males have shown better survival curves against *Pseudomonas entomophila* infection as well as a better bacterial clearance ability against *Providencia rettgeri* than virgin ones, showing absence of reproduction-immunity trade-off [6].

This sexual dimorphism has been also observed in the pathogenesis of some infectious diseases in humans, including tuberculosis [52] which has been the leading cause of death due to a single pathogen (*Mycobacterium tuberculosis*, Mtb), until being surpassed by the COVID-19 pandemic on 2020 [53]. *Mycobacterium marinum* is the most common pathogenic mycobacteria used for infections in non-mammal animal models because it is a natural pathogen of ectotherms causing a granulomatous infection that highly resembles TB in humans [54–56]. The infection of this mycobacterium in the *D. melanogaster* model has also been well characterized. The microorganism is able to kill flies even with very low initial doses, at the early stages of the infection, as it is able to replicate inside *Drosophila* hemocytes [57]. It has also been described that the formation of lipid droplets inside phagocytic cells derived from the production of Upd3 in these cells benefits the intracellular growth of the pathogen, thus resembling the formation of foamy macrophages in humans. As the infection evolves, *M. marinum* causes a progressive loss in energy storage inducing a cachexia-like process that eventually, accompanied by widespread tissue damage, is responsible for killing the flies. This wasting process is mediated by the disruption of the IIS pathway [58].

In this study, we have assessed the role of sexual dimorphism and reproductive status (virgin and mated kept alone or together) on the response to the infection by *M. marinum* in *D. melanogaster*. We have also assessed the differences that these features have on the innate immune response triggered by the pathogen by measuring the expression of Toll- and Imd-dependent AMPs, *drosomycin* and *diptericin* respectively. In addition, due to the importance that metabolism has on the *M. marinum* infection in flies [58], we have assessed the expression levels after the infection of *upd3* and *impl2* based on previous studies that described the role that these molecules play in the metabolic regulation of the host during infections [21,59].

## 2. Materials and methods

### 2.1. Fly stocks and husbandry

Oregon-R-C wildtype flies were obtained from the Bloomington Drosophila Stock Centre (BDSC, Indiana University) and they belong to stock number #5. Flies were raised on a standard cornmeal medium at 25ºC, 65-70% humidity with a 12h light/dark cycle. Male and female flies were aged for 3-5 days before experimentation.

### 2.2. Mycobacterial stock preparation and infection

*Mycobacterium marinum* E11 strain resistant to kanamycin (a kind gift from Wilbert Bitter, Vrije Universiteit Amsterdam) was used [60]. The mycobacterial strain was cultured in 7H9 complete medium complemented with 20µg/ml of antibiotic and placed at 30ºC with constant agitation (170rpm) for 10 days (until an OD_600nm_ of 1.5). The cultures were centrifuged for 5min at 4000g, resuspended in phosphate buffered saline (PBS) with 0.2% Tween 80 and centrifuged again for 5min at 500g to remove clumps. Supernatants were transferred to a new tub, centrifuged for 5min at 4000g and resuspended in 1ml of 7H9 with 15% glycerol. Mycobacterial cultures were then aliquoted and frozen at -80ºC. Each stock was tittered after being frozen at least overnight.

For systemic infections, an aliquot was defrosted and centrifuged at 15000g for 5min. Pellet was rinsed twice with sterile PBS and diluted to the proper concentration. Fourteen nanolitres of the diluted mycobacterial solution were injected systemically employing a nano-injector (Nanoject II, Drummond) into the abdomen of anaesthetized flies.

### 2.3. Experimental design

Flies were divided into three experimental groups: virgin flies, mated flies that were separated by sex after the infection and flies that were allowed to mate throughout the infection (actively mating). Virgin flies were collected at the 2-3h post-eclosion and kept in same-sex groups before and after experimentation. In all experiments, male and female flies were kept in groups of 30 individuals, either only one sex or equally distributed between males and females.

Overall, the experiment consisted of 30 male and 30 female flies per experimental group and 3 biological replicates were performed, with a total of 90 males and 90 females per group. Survival was checked daily and the bacterial load upon death (BLUD) was measured. Dead flies were collected every day, washed with 70% ethanol and rinsed twice with PBS. Each fly was then mechanically homogenized into 200µl of sterile PBS, diluted and plated into 7H10 plates complemented with kanamycin, and incubated for 10 days at 30ºC.

### 2.4. Tolerance and resistance

Previous studies have defined three defensive strategies against infections: qualitative resistance, as the ability to remain uninfected upon pathogen exposure; quantitative resistance, as the ability the reduce the pathogen load; and tolerance, as the ability of the host to maintain the fitness given a certain pathogen load [61,62]. Given our model of study in which the minimum lethal dose of the pathogen is 1, meaning that a single mycobacterium is able to kill the host, we did not assess qualitative resistance but focused on tolerance and quantitative resistance.

Tolerance was defined by the slope of the regression between the host fitness against the pathogen burden [62–67], with more tolerant groups presenting less steep slopes. We used flies’ survival rates as an indicator of host fitness. Uninfected flies were included in the analysis to account for the fitness of each group in the absence of infection, commonly referred to as general vigour [62]. On the other hand, resistance was measured as the Y-intercept of the regression line between the bacterial load upon death (BLUD) and the inoculation dose. When slopes are equal, the lower the Y-intercept the more resistant the group is [62,64,65].

Although the best way to measure these traits would be by evaluating the specific range of bacterial load in the host during the infection [65], in our model is not possible to recover the bacillary load of live flies. Thus, we used the inoculation dose as a controlled measure of the pathogen burden because it highly correlates with both survival times and bacterial load upon death.

### 2.5. Gene expression by quantitative real-time PCR

The differential gene expression was assessed at 24 hours, 5 days and 10 days post-infection. Overall, a total of 9 pools of 3 flies each were analysed per sex, group and time point, obtained from the three independent experiments. All flies were preserved in RNA later (Fisher Scientific, S.L.) and stored at -80ºC before RNA extraction. Total RNA was extracted from flies using the MasterPure™ Complete DNA and RNA Purification Kit (Lucigen) and converted to cDNA with the PrimeScript RT Master Mix (Takara), following the instructions of the kit’s manufacturers for both procedures. cDNA was stored at -20ºC until used.

Quantitative real-time PCR (rt-qPCR) was carried out using the KAPA SYBR® FAST Mix (Sigma) on a LightCycler 480 (Roche Diagnostics) system. RT-qPCR conditions were 95ºC for 5min followed by 40 cycles of 95ºC for 10s and 60ºC or 62ºC for 20s. The specificity of each pair of primers was checked by melting curve analysis (95ºC for 5s, 65ºC for 1min and a continuous rise in temperature to 97ºC at 2.5ºC/s ramp rate followed by 97ºC for 30s). To check reproducibility, each assay was performed with technical triplicates for each biological sample. The relative transcripts levels of target genes were calculated using the 2^-ΔΔCT^ method [68] with r*pl32* used as the reference gene for normalization of target gene abundance.

### 2.6. Statistical analysis

Tolerance and resistance analyses were carried out in GraphPad Prism (version 9.0.0). Survival curves were analysed using Log-rank (Mantel-Cox) test. CFU counts and gene expression data were checked for normality with the Shapiro-Wilk normality test and analysed using one-way ANOVA for multiple comparisons, t-test for single comparisons with normally distributed groups and Mann-Whitney test for non-normally distributed groups. The principal component analysis (PCA) was performed with the Factoextra package in R (version 4.0.1). A linear regression model was used to analyse the tolerance and resistance. Slopes and Y-intercepts were analysed for statistically significant differences with the Extra sum-of-squares F test. Significant differences are as follow: *p≤0.05; **p≤0.01; ***p≤0.001 and ****p≤0.0001 (same for #).

## 3. Results

### 3.1. The effect of reproduction in tolerance and resistance to *M. marinum* infections is sex-dependent

Our study revealed no significant differences in the general vigour of males, independently of their reproductive status. Instead, results showed that the main difference was due to males being reproductively active, rather than having previously mated or not. Thus, males kept together with females were less tolerant but more resistant to the infection by *M. marinum* compared to males kept alone and virgin males (Figure 1a – Table 2).

**Figure 1.**
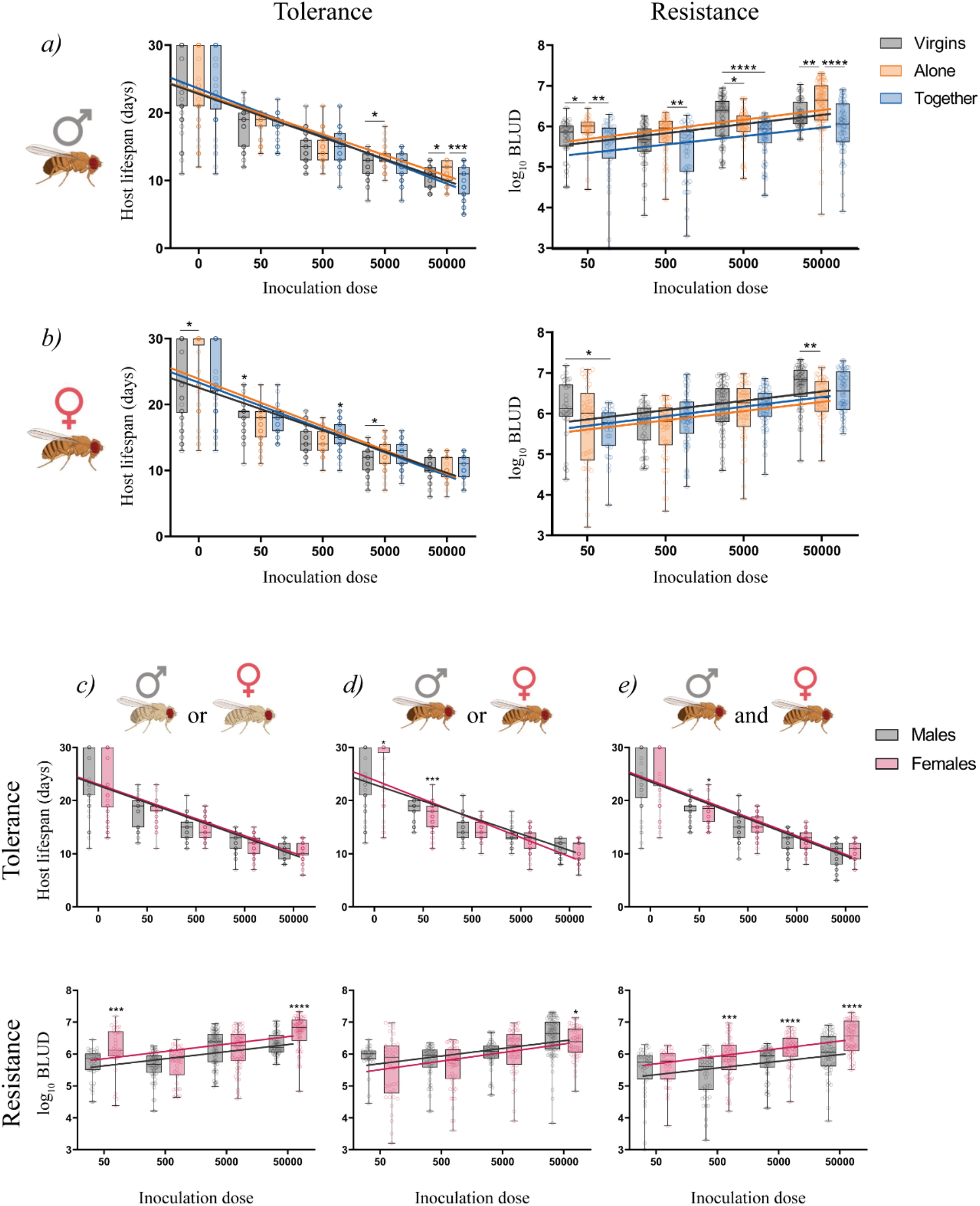
Tolerance and resistance of *D. melanogaster* to *M. marinum* infections depending on the reproductive status or the sex of the flies. (*a*) Males that were not in presence of females (virgins and alone) were more tolerant but less resistant than those kept together with females. (*b*) In females, virgins were more tolerant but less resistant than mated females, independently of the presence or absence of males. When comparing both sexes for each reproductive status, we observed that males were more resistant when virgins or together with females, although no differences in tolerance were found in these groups (*c* and *e*). When flies were kept alone after infection, females showed higher general vigour and resistance, but lower tolerance to infection (*d*). Lines represent the regression lines fitted for each group and each circle represents an individual. Both survival and bacillary load between the groups were analysed independently for each inoculation dose and were tested for normality. Statistically significant differences were represented as follow: *p≤0.05, **p≤0.01, ***p≤0.001, ****p≤0.0001 (Kruskal-Wallis test).

**Table 1.**
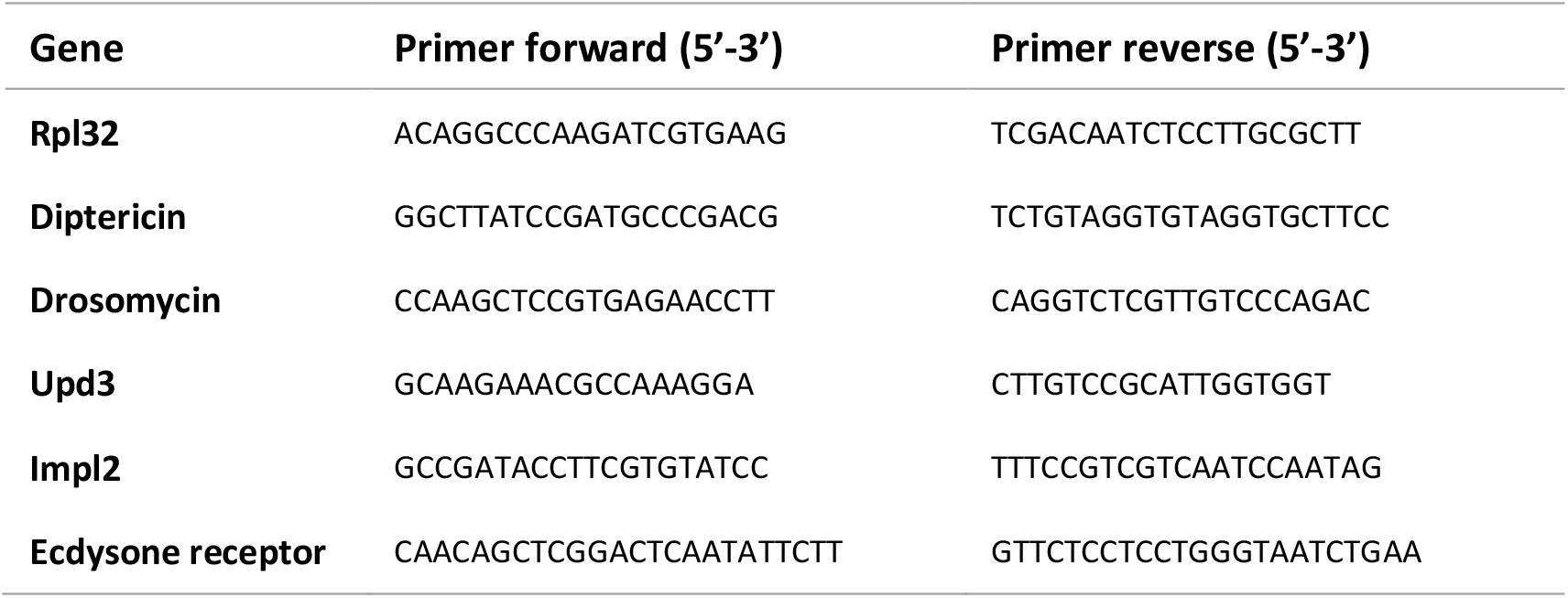
Oligonucleotide sequences used for Real-Time PCR

**Table 2.** Analysis of tolerance and resistance to *M. marinum* infections depending on the reproductive status. Significance (p<0.05) is indicated by asterisks. Slopes for resistance were not statistically significant

| Males |  |  |  |
| --- | --- | --- | --- |
| Tolerance |  |  |  |
| Group | Slope | Comparison | p value |
| Virgins | -3.149 | Alone | 0.3492 |
| Alone | -3.014 | Together | 0.0026** |
| Together | -3.447 | Virgins | 0.0447* |
| Resistance |  |  |  |
| Group | Y-inter | Comparison | p value |
| Virgins | 5.138 | Alone | 0.0325* |
| Alone | 5.196 | Together | <0.0001**** |
| Together | 4.913 | Virgins | <0.0001**** |
| Females |  |  |  |
| Tolerance |  |  |  |
| Group | Slope | Comparison | p value |
| Virgins | -3.165 | Alone | 0.0175* |
| Alone | -3.536 | Together | 0.6273 |
| Together | -3.465 | Virgins | 0.0484* |
| Resistance |  |  |  |
| Group | Y-inter | Comparison | p value |
| Virgins | 5.366 | Alone | <0.0001**** |
| Alone | 4.947 | Together | 0.2354 |
| Together | 5.100 | Virgins | 0.0067* |

On the other hand, the general vigour was significantly lower in virgin females, when compared with both mated groups. In addition, results suggested that females’ response to infection was more conditioned by whether they have mated or not, regardless of whether they were reproductively active at the time of infection or not. Thus, virgin females were more tolerant but less resistant to the infection, compared to both mated groups (Figure 1b – Table 2).

When we compared tolerance and resistance of males and females for each reproductive status independently, data showed that virgin flies differed neither in general vigour nor in overall tolerance to the infection, but males were significantly more resistant (Figure 1c – Table 3). The same pattern was observed in flies that were kept together in equal proportions throughout the procedure: both males and females had the same general vigour and the same tolerance levels, but males showed up to be more resistant (Figure 1e – Table 3). On the contrary, when flies had mated but were kept separated by sex after the infection, females showed increased general vigour and resistance, but reduced tolerance to the infection with *M. marinum* (Figure 1d – Table 3).

**Table 3.**
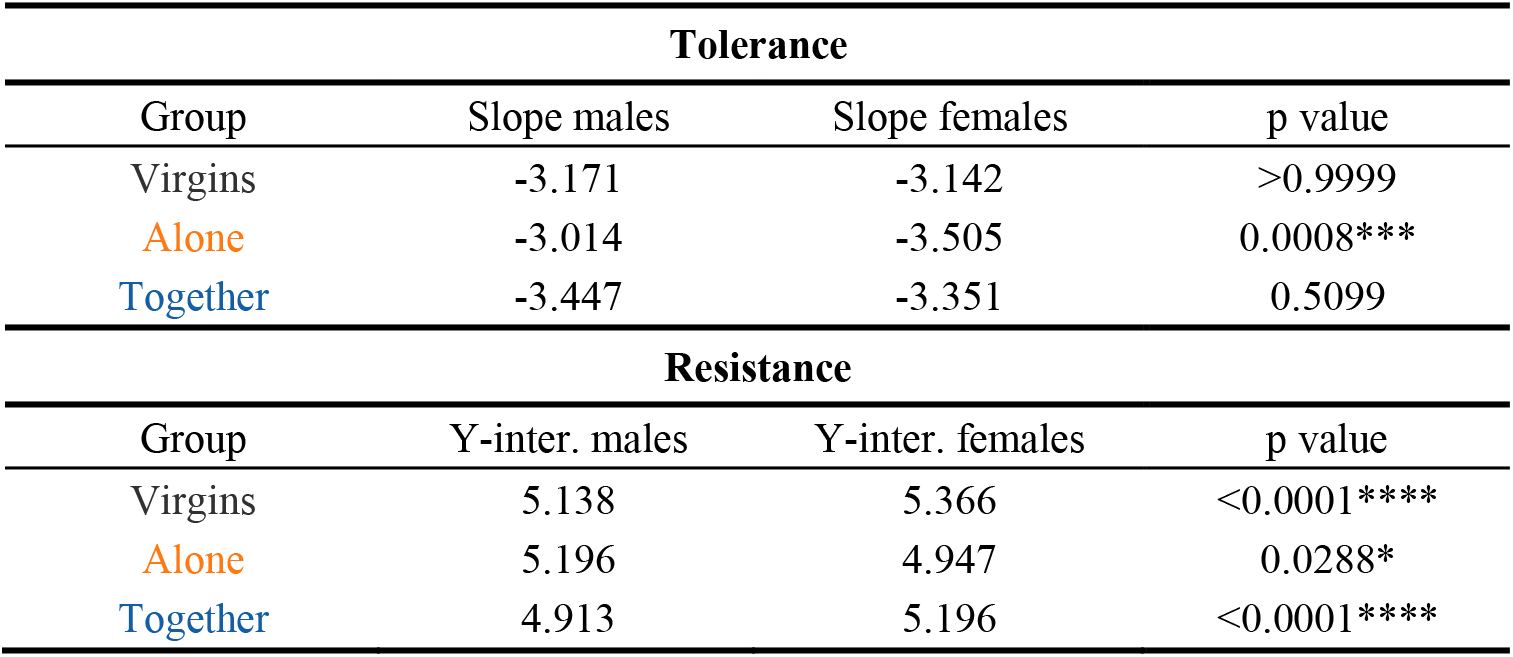
Analysis of tolerance and resistance to *M. marinum* infections depending on the sex. Significance (p<0.05) is indicated by asterisks. Slopes for resistance were not statistically significant.

### 3.2. The humoral innate immune response against *M. marinum* infection depends on the reproductive status of the host

The differences in the innate immune response triggered by the infection in each reproductive status were measuring the expression of the antimicrobial peptides Diptericin and Drosomycin in flies infected with 500 CFU of *M. marinum* at different time points (Figure 2a). These AMPs are regulated by the Imd and the Toll pathways, respectively. In males, only those kept together with females showed immediate significant production of Diptericin, while the other groups showed a delayed production of this AMP. Moreover, virgin males presented significantly increased production of Drosomycin immediately after the infection, while in males alone this production was delayed, and no significant changes were observed in males kept together with females. In females, we observed similar expression profiles as males for each reproductive status: females kept together with males had immediate production of Diptericin, but not Drosomycin; virgin females had delayed production of Diptericin, but early production of Drosomycin; females alone showed delayed or null production of both AMPs.

**Figure 2.**
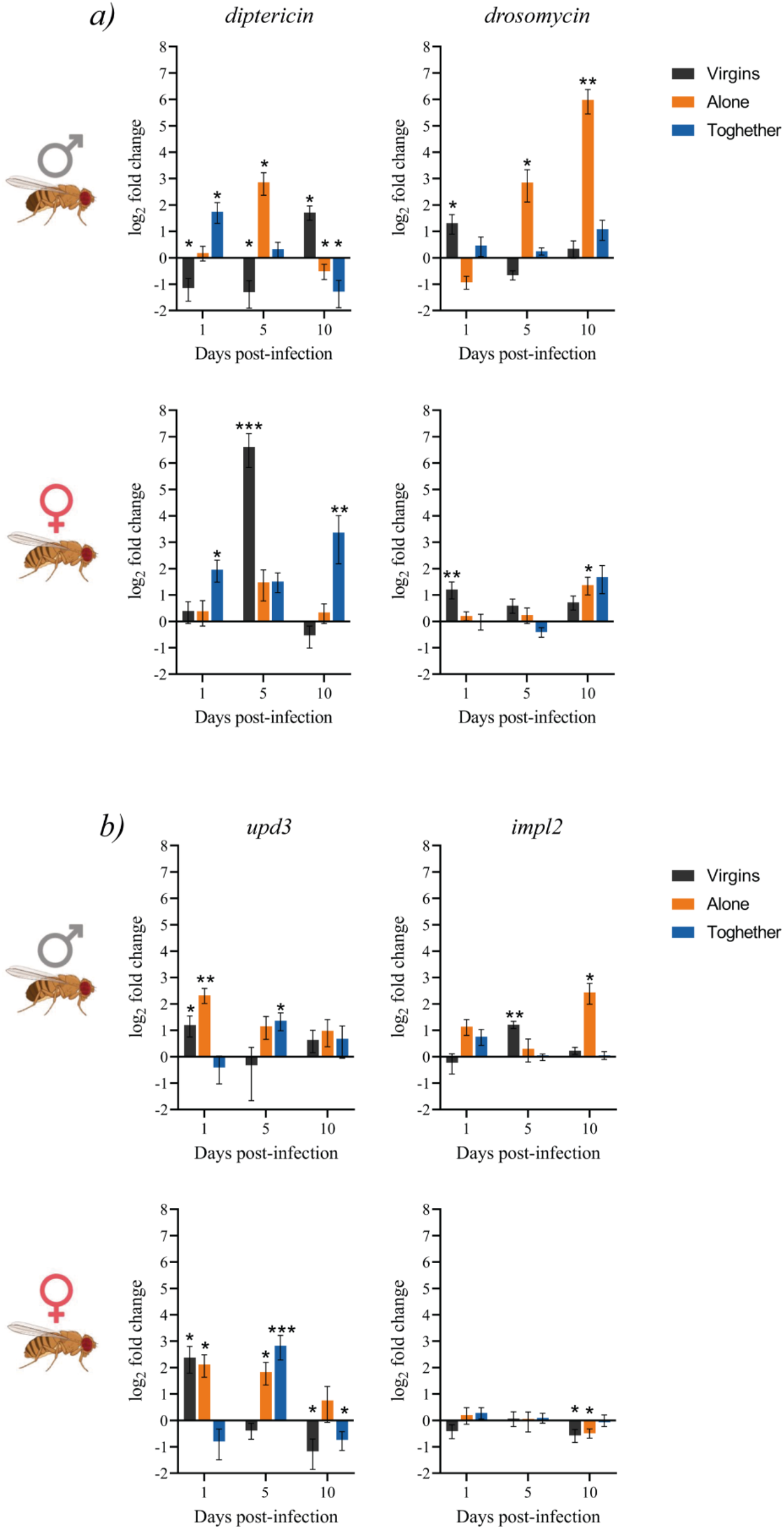
Expression of innate immunity (*a*) and metabolic (*b*) genes during the infection in males and females depending on their reproductive status. Gene expression relative to the internal control gene *rpl32* was quantified in 9 replicate pools of 3 males or females exposed to the infection with *M. marinum* relative to their expression in uninfected controls. Each group at each time-point was compared to its relative control independently (the line in each graph represents the controls’ relative expression with a log_2_ fold change of 0). Data was analysed for normality and significant differences were represented as follow: *p≤0.05, **p≤0.01 (Welch’s corrections for normally distributed data and Mann-Whitney test for not normally distributed data).

These results for the expression levels showed correlation with the phenotypic results obtained previously. We found that the activation of the Toll pathway (production of Drosomycin) correlated with more tolerant but less resistant phenotypes in both males (virgins and alone) and females (virgins). In addition, the more sustained was this activation (males alone) the lower resistance was observed. On the other hand, we also observed that while early activation of the Imd pathway correlated with the more resistant groups (males and females together), late or null activation of this pathway seemed to be linked with reduced resistance with the exception of females alone. These correlations were also observed when comparing the expression levels between males and females for each reproductive status (Supp. Figure 1a). We found that the less resistant groups (virgin females, females kept together with males, and males alone) showed significantly increased production of both AMPs late in the infection, while the more tolerant group (males alone) had increased production of Drosomycin. Finally, results also suggest that females kept alone after mating may acquire their resistance through a mechanism that does not involve the humoral innate immunity, as they did not show any activation of the Imd pathway throughout the infection.

### 3.3. The metabolic regulation during infection is altered by the reproductive status of the host

The link with the metabolism, especially with the insulin/insulin-like signalling pathway (IIS), together with the suggested role for Upd3 and Impl2 as “selfish immune factors” (SIFs), prompted us to investigate whether a differential expression of these molecules within the different reproductive status of the host might also influence the tolerance and resistance levels to the infection by *M. marinum* (Figure 2b).

In males, both virgins and those kept alone showed significant early overexpression of *upd3*, which was not sustained later in the infection. On the other hand, those males that were kept together with females during the whole procedure presented a significant overexpression of this gene later in the infection. In females, virgins showed a significant early but not sustained overexpression of *upd3*, while females that have mated presented a significantly increased expression of this gene later in the infection whether they were in the presence or absence of males. Overall, these results suggest a dual role for *upd3*, as its early expression correlated with more tolerant but less resistant groups while its expression later in the infection correlated with more resistant and less tolerant groups.

Finally, assessing the expression of *impl2* during the *M. marinum* infection we observed no significant differences among females depending on their reproductive status. On the other hand, both virgin males and males alone showed a significant increase in the expression of this gene later in the infection, but not sustained. These results suggested that, in males, the punctual expression of *impl2* might increase tolerance to the infection with *M. marinum*, but not in females.

We also compared the expression levels of both genes between males and females for each reproductive status independently (Supp. Figure 1b). We found that females have an overall increased expression of *upd3* compared to males, while *impl2* seemed to only play a role in males with its late production correlating with increased tolerance (males alone).

### 3.4. Induction of the ecdysone receptor correlated with basal levels prior to the infection in males, but not in females

We also evaluated the expression levels of the ecdysone receptor (EcR) due to the tight relationship of the Ecdysone pathway with immunity and reproduction. We measured the relative expression of this gene after the infection and the basal expression levels when injected with PBS for each reproductive status (Figure 3c).

**Figure 3.**
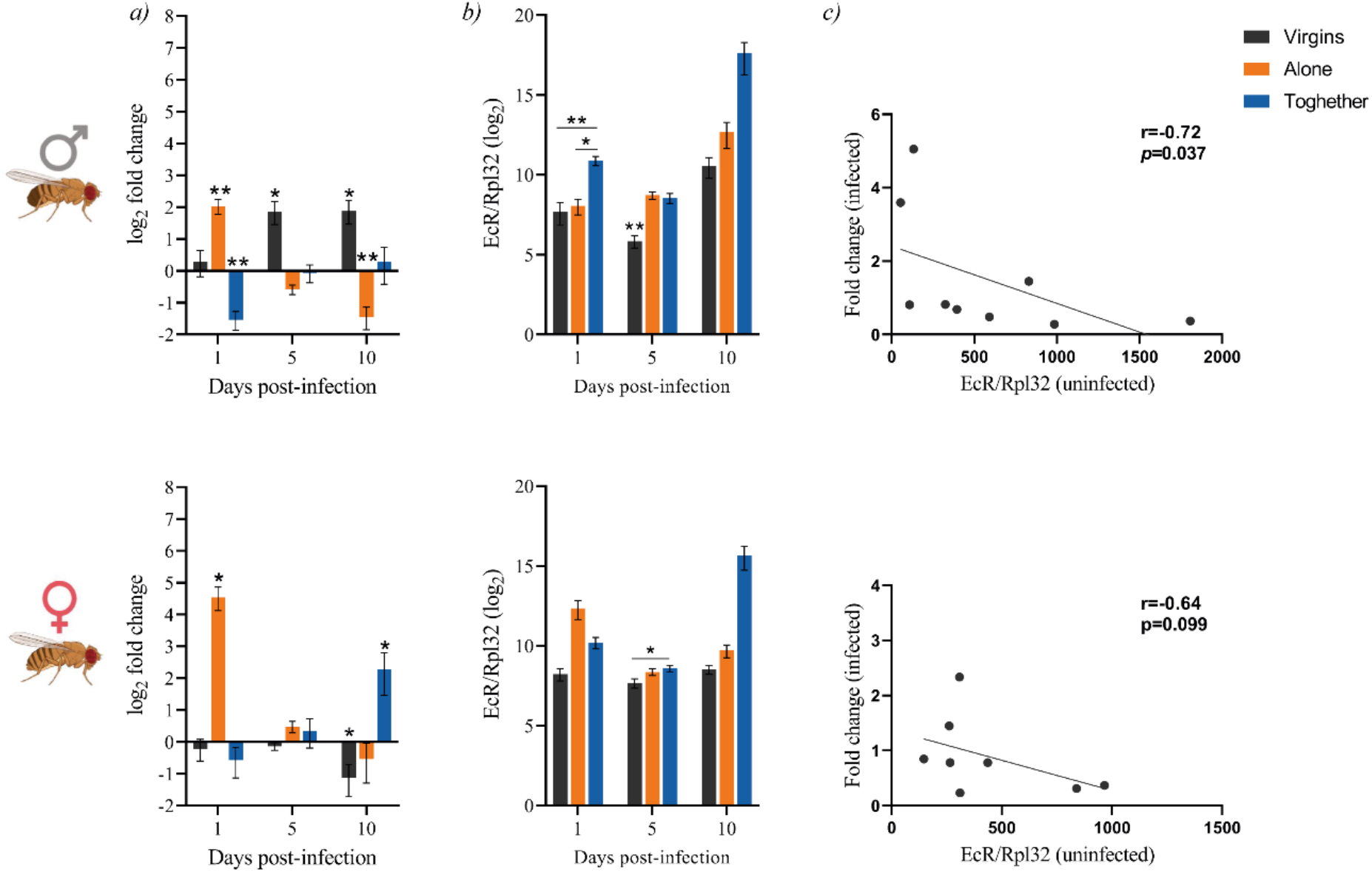
Expression of the ecdysone receptor (EcR) in males (up) and females (down) depending on their reproductive status after the infection (*a*). Gene expression relative to the internal control gene *rpl32* was quantified in 9 replicate pools of 3 males and 3 females each exposed to the infection with *M. marinum* relative to their expression in uninfected controls. Each group at each time-point was compared to its relative control independently (the line in each graph represents the controls’ relative expression with a log_2_ fold change of 0). Data was analysed for normality and significant differences were represented as follow: *p≤0.05, **p≤0.01 (Welch’s corrections for normally distributed data and Mann-Whitney test for not normally distributed data). Basal expression levels of the EcR gene in uninfected flies (*b*) were calculated using the 2^-ΔCT^ method with the *rpl32* gene for normalization (all values were multiplied by 10^4^ for more visual results). Groups were compared independently for each time-point. Data was analysed for normality and significant differences were represented as follow: *p≤0.05, **p≤0.01 (Kruskal-Wallis test). Finally, the correlation between the basal levels and the fold change after infection was performed using the nonparametric spearman correlation test (*c*).

We found that virgin males significantly induced the expression of the receptor later in the infection, while males alone did it immediately after. On the other hand, the infection triggered a significant repression of the receptor in those males kept together with females (Figure 3a). We also found that males presented a significant correlation between the basal level of EcR and the induction of this receptor post-infection, in which lower basal levels translated into higher activation of this gene post-infection (Figure 3b and 3c). In females, virgins showed no changes neither in the basal levels nor in the induction of EcR after the infection. Females that had mated showed similar patterns as their males counterparts, while females kept alone showed decreasing induction of EcR when infected, but no significant changes on the basal levels (Figure 3a and 3b). However, the correlation between basal levels and induction of EcR after the infection was not statistically significant in females (Figure 3c).

The results also showed a link between the basal expression levels of EcR and the resistance to the infection. Those groups with higher basal expression levels (males and females kept together and females alone) were more resistant to the infection, while those groups with lower basal levels (virgin flies and males alone) were less resistant.

When comparing both basal and induction levels between males and females (Supp. Figure 2a), we observed that virgin males presented lower basal levels but increased production of EcR after the infection, while flies separated after the infection showed no significant different basal levels but females increased its production after the infection. Finally, when flies were kept together throughout the procedure, no differences at basal levels among sexes were found, but a late induction in females.

### 3.5. Gene expression of *D. melanogaster* upon infection shows a marked sexual dimorphism marked by Impl2 and Upd3 in males and females, respectively

Finally, we assessed the overall correlation between all the genes studied. The principal component analysis (PCA) (Figure 4a) performed with the relative expression levels of infected flies including all time-points for each condition revealed that, in males, all three groups presented significantly different gene expression profiles, which linked with all three groups presenting different tolerance and resistance profiles as well (Figure 4b). In females, only virgins presented a significantly different gene expression profile (Figure 4b), also linking with the phenotypic results. Finally, the analysis of the variable contribution (Figure 4b) for dimension 1 (PC1), revealed that, in males, this difference was mainly driven by the differential expression of Impl2, together with Drosomycin and Upd2, while in females the main differences were driven by Upd3, together with EcR and Impl2 (Figure 4c).

**Figure 4.**
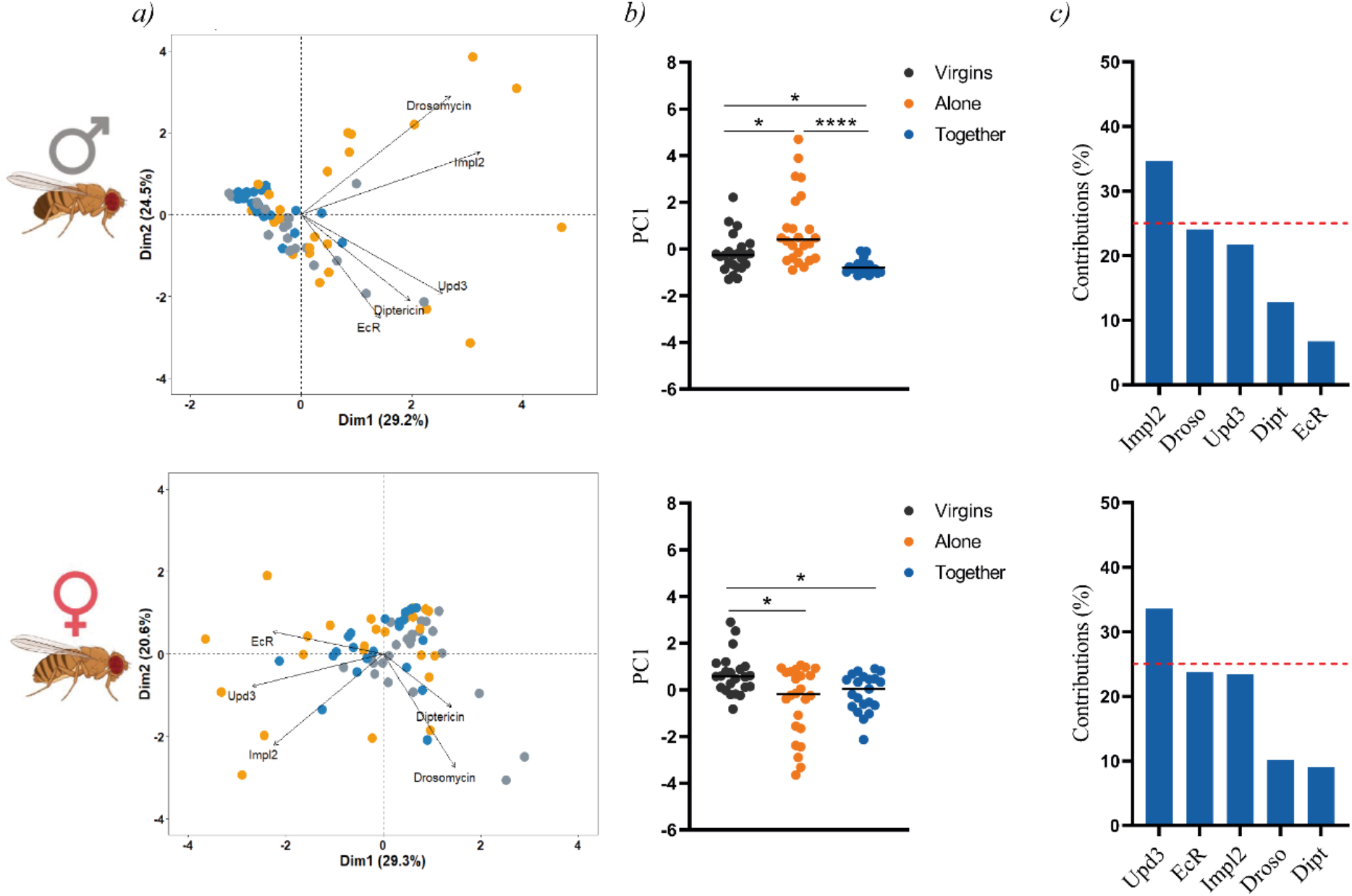
Heterogeneity of gene expression among flies with different reproductive status. Principal component analysis (PCA) based on expression of genes of interest in males and females infected with M. marinum at all time-points (*a*), PC1 scores (*b*) and variable contribution to PC1 (*c*). Each circle represents an individual and lines are means. Statistically significant differences were represented as follow: *p≤0.05, ****p≤0.0001 (Kruskal-Wallis test).

## 4. Discussion and conclusions

Typically, laboratory studies have revealed a significant “cost of mating” to *Drosophila* females in the form of reduced longevity. However, here we present that virgin females show a significant decrease in general vigour compared to mated females when uninfected, which is not observed in virgin males. This phenomenon has only been described previously for wild-caught flies [69]. Previous studies described that both males and females reduce their resistance to some infections after mating [45,50,70]. In contrast, data presented herein shows that the opposite happens in both sexes during infection with *M. marinum*. Males increase their resistance to the infection after mating when they are together with females, although at the expense of reducing their fitness, while females increase their resistance to *M. marinum* infection after mating, independently of the presence or absence of males. Overall, phenotypic results from this study show that the sexual dimorphism observed in *D. melanogaster* in the outcome of *M. marinum* infection is highly related to the reproductive status of the host. This probes that when assessing the differences in the tolerance and/or resistance of *D. melanogaster* to other infections, the hosts’ reproductive situation needs to be characterised.

Regarding the hormonal levels and their relation with the immune response, our data show that certain EcR levels are required for the flies after the infection, as flies with higher basal levels presented lower induction or repression of the gene post-infection. However, this correlation was only statistically significant in males. Previous studies showed that ecdysone induces the expression of the receptor PGRP-LC and, thus, modulates the Imd pathway [39]. Other studies had related the Imd pathway in the control of resistance to infections [71], while its negative regulation mediates tolerance to infections [72], although any of these were performed in mycobacterial infections. Our data support these findings, as those flies with higher basal levels of EcR show higher expression levels of Diptericin when infected and correlate with the more resistant phenotypes, while flies that did not show early production of this AMP were more tolerant to the infection.

Several studies performed with a wide range of pathogens have described the Toll pathway as key in determining resistance to infections [32,73,74], and even to be essential for resistance in males, but not in females [29]. However, no role in increasing tolerance to infection has been described for this pathway previously. In our study, we showed that production of the Toll-dependant AMP, Drosomycin, early in the infection was linked with increased tolerance but reduced resistance to *M. marinum* infection in flies, while their activation in later stages of the infection correlated with increased resistance. Altogether, these data suggest a role for the Toll pathway in determining both tolerance and resistance to *M. marinum* infections. Recent studies have described a Toll-dependent metabolic switch that directs fatty acids from neutral cellular storage toward phospholipid biosynthesis [75], which might reduce the lipid droplet accumulation inside phagocytic cells and, thus, hindered the intracellular replication of *M. marinum*.

This study also suggests a dual role for Upd3 in *M. marinum* infections. Our data correlate with previous studies that linked Upd3-deficient flies with a reduction of lipid droplet accumulation within cells and reduced bacillary loads [59], showing that flies with increased expression of Upd3 early in the infection presented higher bacillary loads. Yet, this study also suggests a second role for Upd3, as its production late in the infection correlate with increased resistance, probably due to its role in increasing the metabolism of phagocytic cells [23]. In addition, our data also suggest that the increased resistance in females alone might be driven by Upd3 rather than by innate immune pathways.

Finally, the expression of Impl2 during *M. marinum* infections seems to be only relevant in males that are not actively mating and to be related to increased bacillary loads, although further studies should be performed to determine its role during mycobacterial infections in *D. melanogaster*.

## Concluding remarks

It has been extensively studied that males and females respond differently to the same infections. Many of these studies have focused on establishing the physiological basis for these differences, but very few have studied how the reproductive status of the host affects males and females and the role it plays in the response to infection. This study aimed to show that the intrinsic differences observed between males and females during the infection by *M. marinum* were tightly related to the reproductive status of the host and that reproduction affects males and females differently and that when assessing the sexual dimorphism of *D. melanogaster*, the hosts’ reproductive situation need to be characterised.

Here we show that being actively mating (kept together) increased the resistance but reduced the tolerance to the infection in males and females. The same phenotype was found in mated females regardless if they were kept alone or together after mating.

The results obtained in this study also suggest a possible role for the Toll pathway in determining tolerance to infection, while it appears that the IMD pathway is associated with increased resistance. Results also showed a correlation between basal levels of EcR and its induction after the infection, suggesting that a certain amount of this receptor is required upon infection. This phenomenon was observed in both males and females in all reproductive statuses, although only in males was statistically significant. In addition, basal levels of EcR also correlated with higher expression levels of Diptericin.

Finally, we have also proposed a dual role for Upd3 upon infection with *M. marinum*. Our results show that an early production of this molecule correlate with higher bacillary loads. This might be linked with the Upd3-mediated lipid droplets accumulation inside phagocytic cells, which favours mycobacterial replication. However, the late production of this molecule is associated with lower bacillary loads, most likely due to its role in increasing phagocytic cells metabolism. Finally, we could only validate the role for Impl2 in increasing the tolerance in non-virgin non-actively mating males.

## 5. Acknowledgements

This research has been funded by “La Caixa” Foundation (ID 100010434), under agreement LCF/PR/GN16/10290002, by Spanish Government-FEDER Funds through PI20/01431 grant, and by the “CIBER Enfermedades Respiratorias” Network (CIBERES).

