## Supplemental Figures for "The reproductive status of the host determines the tolerance and resistance to *Mycobacterium marinum* infection in *Drosophila melanogaster*"

### SUPPLEMENTARY FILE

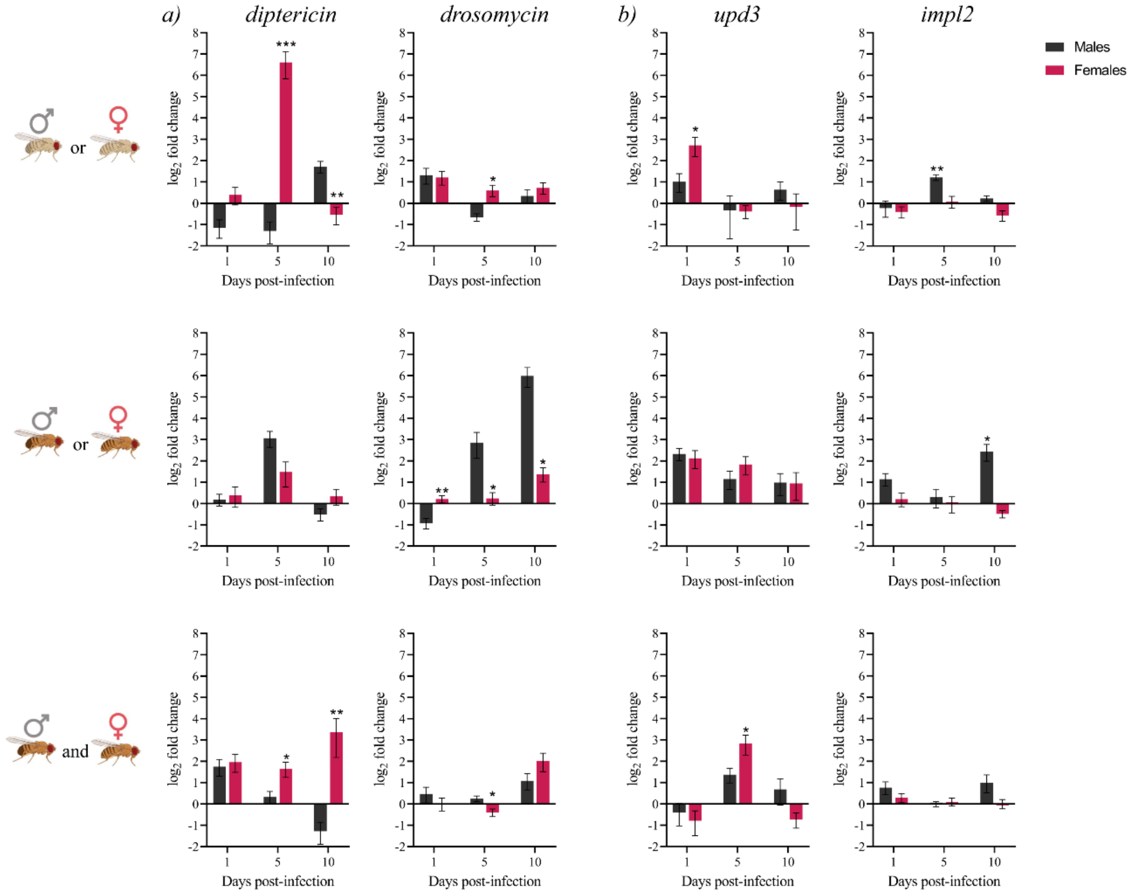


**Supplementary Figure 1|** Expression of innate immune (*a*) and metabolic (*b*) genes during the infection in flies depending on their sex for each reproductive status independently. Gene expression relative to the internal control gene *rpl32* was quantified in 9 replicate pools of 3 males or females exposed to the infection with *M. marinum* relative to their expression in uninfected controls. Each time-point was compared independently (the line in each graph represents the controls' relative expression with a $\log_2$ fold change of 0). Data was analysed for normality and significant differences were represented as follow: *p≤0.05, **p≤0.01, ***≤0.001 (Welch's corrections for normally distributed data and Mann-Whitney test for not normally distributed data).

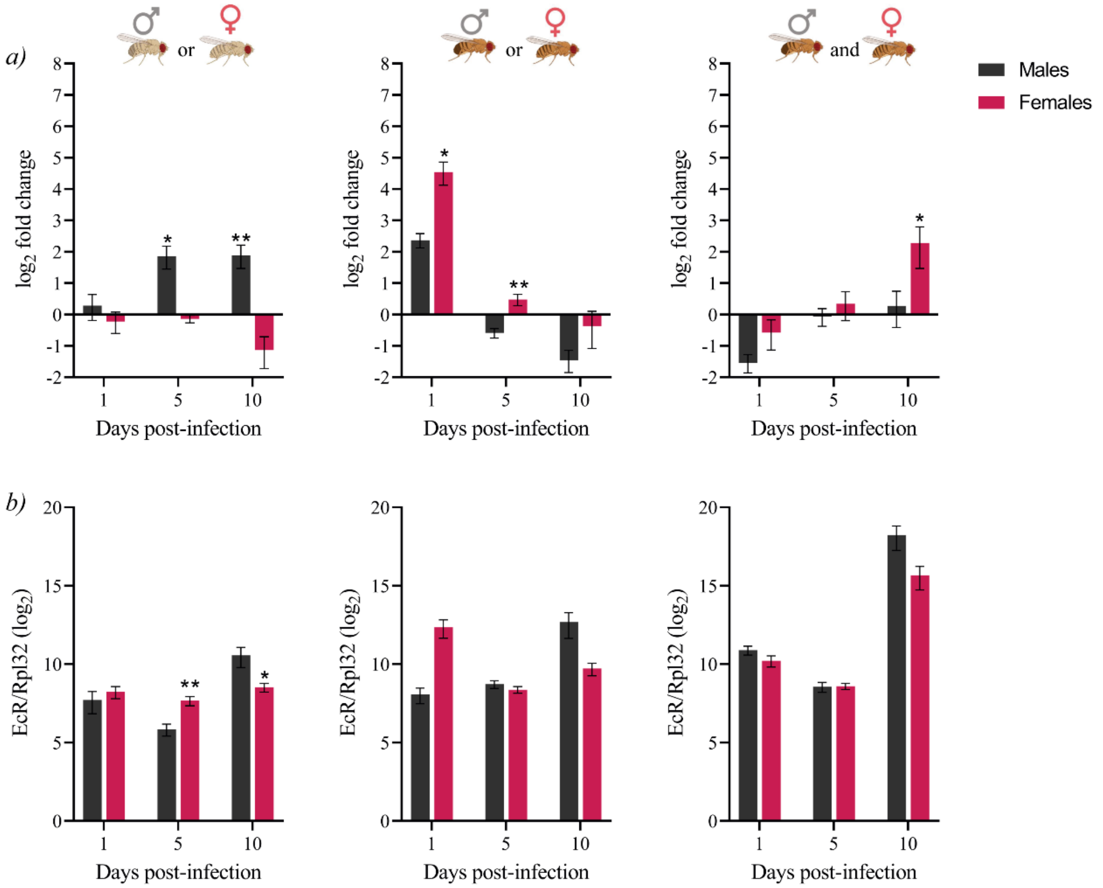


**Supplementary Figure 2|** Expression of the ecdysone receptor (EcR) in flies depending on their sex for each reproductive status independently ***(a)*.** Gene expression relative to the internal control gene *rpl32* was quantified in 9 replicate pools of 3 males and 3 females each exposed to the infection with *M. marinum* relative to their expression in uninfected controls. Each time-point was compared independently (the line in each graph represents the controls' relative expression with a $log_2$ fold change of 0). Data was analysed for normality and significant differences were represented as follow: *p≤0.05, **p≤0.01 (Welch's corrections for normally distributed data and Mann-Whitney test for not normally distributed data). Basal expression levels of the EcR gene in uninfected flies ***(b)*** were calculated using the $2^{-\Delta CT}$ method with the *rpl32* gene for normalization (all values were multiplied by $10^4$ for more visual results). Groups were compared independently for each time-point. Data was analysed for normality and significant differences were represented as follow: *p≤0.05, **p≤0.01 (Kruskal-Wallis test).
